# Classification of RNA-Seq Data via Gaussian Copulas

**DOI:** 10.1101/116046

**Authors:** Qingyang Zhang

## Abstract

RNA-sequencing (RNA-Seq) has become a preferred option to quantify gene expression, because it is more accurate and reliable than microarrays. In RNA-Seq experiments, the expression level of a gene is measured by the count of short reads that are mapped to the gene region. Although some normal-based statistical methods may also be applied to log-transformed read counts, they are not ideal for directly modeling RNA-Seq data. Two discrete distributions, Poisson distribution and negative binomial distribution, have been commonly used in the literature to model RNA-Seq data, where the latter is a natural extension of the former with allowance of overdispersion. Due to the technical difficulty in modeling correlated counts, most existing classifiers based on discrete distributions assume that genes are independent of each other. However, as we show in this paper, the independence assumption may cause non-ignorable bias in estimating the discriminant score, making the classification inaccurate. To this end, we drop the independence assumption and explicitly model the dependence between genes using Gaussian copula. We apply a Bayesian approach to estimate covariance matrix and the overdispersion parameter in negative binomial distribution. Both synthetic data and real data are used to demonstrate the advantages of our model.

## 1 Introduction

RNA-sequencing (RNA-Seq) is a revolutionary tool for the study of transcriptomes (Mardis [2008]; Wang et al. [2009]). Compared to hybridization-based microarrays, RNA-Seq eliminates the need for species-specific sequence information and provides more reliable measurements for gene expression (Marioni et al. [2008]). The huge number of reads produced by RNA-Seq experiment enables researchers to better detect novel transcripts and quantify the gene expression in ultra-high resolution. Essentially, RNA-Seq consists of three distinct phases: (1) RNA is isolated from tissue and segmented to an average length of 200 base pairs; (2) RNA segments are reverse transcribed to cDNAs; (3) The cDNAs are mapped to reference transcriptome or genome. An RNA-Seq experiment usually produces tens of millions of short reads between 25 and 300 base pairs in length. The number of reads mapped to each transcript provides a digital measure of transcript abundance.

Poisson distribution and negative binomial distribution are commonly used distributions to model RNA-Seq data. Based on these two distributions, numerous methods have been proposed to detect the differentially expressed genes, including but not limited to edgeR (Robinson & Smyth [2008]), DESeq2 (Love et al. [2014]), baySeq (Hardcastle & Kelly [2014]), BBSeq (Zhou et al. [2011]), and SAMseq (Li & Tibshirani [2013]). Despite the significant advances in differential expression analysis, the progress on classification of RNA-Seq data is relatively recent. Witten (Witten [2011]) developed a Poisson linear discriminant analysis (PLDA) by assuming that the data follow a Poisson distribution. However, in the presence of overdispersion, the Poisson assumption might not be appropriate. Dong et al. (Dong et al. [2016]) further extended the Poisson classifier to a negative binomial classifier (NBLDA), and explored how the dispersion affects the classifications. Other classifiers developed for RNA-Seq data include logistic regression model and partial least square method (Tan et al. [2014]). Due to the difficulty of modeling correlated counts, most existing classifiers assume that all genes are independent of each other. However, as pointed out by Dong et al. (Dong et al. [2016]), this assumption is very restrictive and may not be realistic in practice. The objective of the paper is to numerically assess the effect of independence assumption on classification of RNA-Seq data, and to develop a new classifier incorporating the dependence between genes using continuous latent variables and Gaussian copula. A Metropolis-Hasting algorithm in combination with Gibbs sampler (Lee [2014], Liu & Daniels [2006]) is adopted to estimate the covariance matrix and overdispersion parameters in our model. Our new classifier explicitly models two important aspects of RNA-Seq data: overdispersion of read counts and correlation between genes, therefore provides accurate parameter estimate and sample classification.

Copula is an important tool in modeling the dependence between random variables of any type. It is especially useful for modeling multiple discrete variables whose joint distribution can be extremely complicated. To begin with, we provide a short review of copula function and Gaussian copula. Consider a vector of random variables (*X*_1_, *X*_2_,…, *X_p_*), the copula function of (*X*_1_, *X*_2_,…, *X_p_*), *C*: [0,1]^*p*^ → [0,1], is defined as the cumulative distribution function (cdf) of (*F*(*X*_1_),*F*(*X*_2_),…,*F*(*X_p_*)):

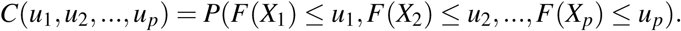

By definition, a copula function is a multivariate distribution function where the marginal of each random variable is uniform. Sklar’s Theorem guarantees that any multivariate distribution can be expressed with univariate marginals and a copula function which links the marginals. In practice, we can completely separate the choice of marginals and the choice of copula. Popular copulas include, but not limited to Gaussian copula, Student’s t copula, Clayton’s copula and Frank’s copula (Nelson [1999]). Clayton’s copula and Frank’s copula both belong to the bivariate Archimedean copula family. For more than two dimensions, Gaussian copula is convenient to model the complex correlation structure (both positive and negative correlations). The Gaussian copula is based on multivariate normal distribution:

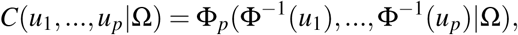

where Φ represents the cdf of standard normal distribution, Φ_*p*_(…|Ω) represents the cdf of p-dimension normal distribution with correlation matrix Ω. To connect discrete marginals and continuous copula function, we introduce a latent variable and treat the observed count as the discretized value of the continuous latent variable (in spirit, it is same as multivariate Probit model).

The remainder of this paper is structured as follows. In Section 2, we formally describe the statistical framework for classification of RNA-Seq data and introduce a Bayesian approach to estimate unknown parameters. Numerical studies are conducted to compare different models and classifiers in Section 3. In Section 4, we apply the proposed method to two real data sets including the cervical cancer data and HapMap data. We discuss and conclude this paper in Sections 5 and 6.

## 2 Methods

In this section, we propose a new classifier for RNA-Seq data based on copula function. We assume that the data follow a complex multivariate distribution with negative binomial marginals. The correlation between genes can be described by a Gaussian copula. A general Bayesian framework for estimating parameters in each class is discussed.

### 2.1 Negative binomial distributions for marginal model

First, we consider only one class. Let ***x =*** (***x***_1_…,***x***_*n*_)^*T*^ be the *n* × *p* data matrix, where ***x***_*ij*_ denotes the observed number of reads mapped to gene *i* in sample *j, i* = 1,2,…, *p* and *j* = 1,2,…,*n*. We consider the following negative binomial distribution *F_ij_*, *i* = 1,2,…,*p*, *j* = 1,…,*n* for marginals (Dong et al. [2016]):

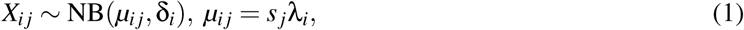

where *μ_ij_* **=** *E* (*X_ij_*), *s_j_* is the size factor to scale read counts for the *j*^th^ sample due to different sequencing depth, *λ_i_* is the total number of reads for gene *i*, and *δ_i_* is the overdispersion parameter for gene *i*, i.e., *V*(*X_ij_*) = *μ_ij_* + *μ_ij_*^2^δ_*i*_. The estimates of *λ_i_* and *s_j_* in (1) are straightforward:

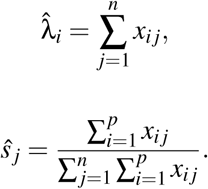

For overdispersion parameter *δ_i_* the moment estimate and shrinkage estimate (Yu et al. [2013]) are commonly used estimates. However, both methods suffer from instability, e.g., the moment method sometimes gives a negative value. In this paper, we treat *δ_i_* as unknown parameter, which is to be estimated jointly with other parameters in a Bayesian framework.

### 2.2 Gaussian copula for dependence between genes

In general, the analysis of correlated counts might be difficult because of the lack of suitable discrete multivariate distribution that can model complex correlation structures. To surmount this difficulty, we model the correlation via Gaussian copula, so that the correlation between read counts can be created through the correlation of the continuous latent variables. Let ***Z***_*j*_ = (*Z*_1*j*_,…,*Z_pj_*)^*T*^ be the Gaussian latent variables (with unit variance) of ***X***_*j*_, and ***Z***_*j*_ ~ *N_p_*(**0**, Ω), where Ω represents the covariance or correlation matrix. The observed counts ***x***_*j*_ are the discretized values of ***z***_*j*_ by quantile matching. The relation between *X_ij_* and *Z_ij_* can be interpreted as follows:

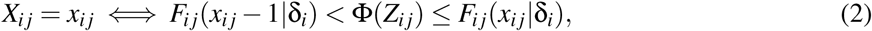

where *F_ij_*() represents the cumulative distribution function of variable *X_ij_*, *x_ij_* takes nonnegative integer values 0,1,2,… and *F_ij_*(–1 |δ_*i*_) = 0 by definition of cdf.

The Gaussian copula of latent variables has the following simple form:

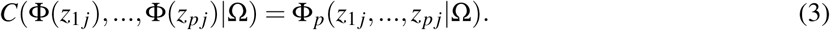

Based on (2) and (3), the likelihood function can be obtained immediately:

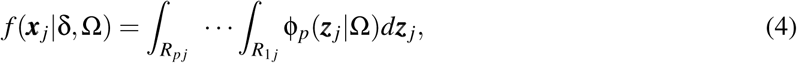

where δ = (δ_1_,…, δ_*p*_), φ_*p*_ denotes the multivariate normal density of dimension *p*, with mean vector **0** and unit marginal variance. The endpoints defining integration region *R_ij_ =* (*L_ij_*, *U_ij_*] are specified as a function of the parameter δ_*i*_

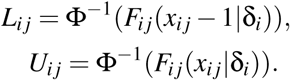

### 2.3 Bayesian estimation of parameters

Finding the maximizer of (4) is impractical due to the complexity and non-convexity of the likelihood function. Here, we consider the Bayesian approach proposed by Lee (Lee [2014]; Liu & Daniels [2006]) for parameter estimate. The posterior distribution of the parameters and latent variables can be written as follow:

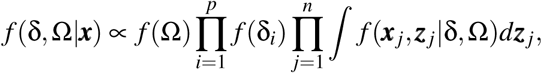

where the priors are specified as *f*(δ_*i*_) *= IG*(α_0_, β_0_), and *f*(Ω) = *IW*_***p***_(Ψ_0_, *ν*_0_). We use *IG*( α_0_, β_0_) to denote the inverse-gamma distribution with shape parameter α_0_ and rate parameter β_0_, and use *IW*(Ψ_0_, *ν*_0_) to denote the inverse-Wishart distribution with scale matrix Ψ_0_ and degree of freedom Ψ_0_. The Gibbs sampling can be implemented based on the following conditional distributions:

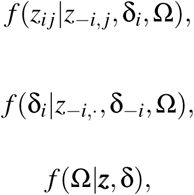

where z_–*i,j*_ = (*z*_1*j*_,…,*z*_(i–1)_*j*,*z*_(*i*+1)_*j*,…,*z_pi_*),*z*_–i_, = {*z*_–i,j_, *j*=1,…,*n*}, δ_–*i*_=(δ_1_,…, δ_*i*–1_, δ_*i*+1_,…, δ_*p*_), ***z*** = (*z*_1_,…,*z_n_*)^*T*^. Conditioning on {*z*_–*i*,*j*_, δ_*i*_, Ω }, latent variable ***Z***_*ij*_ follows a truncated normal distribution, where the mean and variance depend on {*z*_–*i*,*j*_, Ω}, and the endpoints depend on {*x_ij_*, δ_*i*_}. We use a routine Metropolis-Hasting algorithm to sample δ_*i*_ and use a parameter-expanded reparameterization and Metropolis-Hasting algorithm (PXMH, Lee [2014]; Liu & Daniels [2006]) to sample Ω. Details about these algorithms are provided in the Appendix.

### 2.4 Classification

We consider the classification problem when the RNA-Seq were conducted over multiple classes, i.e., *K* ≥ 2. Let *y_j_* ∈ {1,2,…,*K*} denotes the class label of sample *j*, i.e., *y_j_* = *k* ⇔ *j* ∈ *C_k_*. The marginal distribution of *X_ij_* in class *k* can be formulated in a similar way to (1):

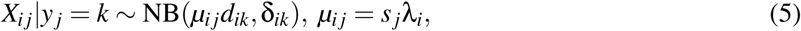

where *d_ik_* and δ_*ik*_ are gene- and class-specific parameters among the K classes. The overdispersion parameter δ_*ik*_ can be estimated in the PXMH algorithm and *d_ik_* can be estimated by

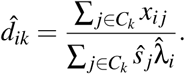

Based on the trained models for all the K classes, the class label for new observation ***x\**** can be predicted. By Bayes’ rule:

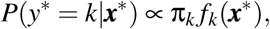

where *f_k_* is the probability density function for class *k*. The prior probability π_*k*_ can be estimated by 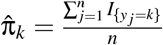. We assign the new observation ***x***\* to class *k* that maximizes the following discriminant score (posterior probability):

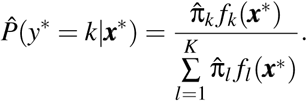

## 3 Simulation study

We conducted two simulation studies with *K* = 2 to benchmark our new classifier. In Simulation I, we evaluated the performance of the Bayesian approach in estimating the discriminant score *P*(*y*\* = 1|***x***\*) under different correlation settings. In Simulation II, we compared the performance of six classifiers under different settings of dispersion and correlation strength.

### 3.1 Simulation I

Given the correlation matrices Ω, *k* = 1,2, we generated data in two steps:

- Step 1: Simulate latent variables *z_jk_* ∼ *N_p_*(**0**,Ω_*k*_), *j* = 1,…,*n_k_,k* = 1,2, where Ω is the correlation matrix or covariance matrix, *n*_1_ = *n*_2_ = 50 and *p* = 50
- Step 2: Transform ***z***_*jk*_ to ***x***_*jk*_, *j* = 1,…,*n* using (2), where *μ_i_*_1_ = 20, *μ*_*i*2_ = *μ_i_*_1_ + Δ_*i*_, Δ_*i*_ ∼ *Unif* (–15,15) and δ_*i*1_ = δ_*i*2_ ∼ *Unif* (1,10) for *i* = 1,…, *p*

We compared the copula-based model with Dong et al.'s independence model under two settings of autoregressive correlation structure Ω_1_(*i*, *j*) = Ω_2_(*i*, *j*) = exp(–*a*|*i* – *j*|): (1)*a* = 0.5; (2)*a* = 1.5. For each class, we trained the model using half of the samples (25 samples randomly chosen in each class) and then calculated the discriminant score *P*(*y*\* = 1|***x***\*) for each of the rest samples. For both models, we estimated *μ_ik_* by 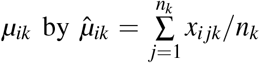. For independence model, we estimated the overdispersion parameter δ_*ik*_ using R package *sSeq,* which implements the shrinkage method by Yu et al. The score *P*(*y*\* = 1 |***x***\*) was then estimated under independence assumption. For the copula-based model, we jointly estimated δ_*ik*_ and Ω_*ik*_ using PXMH algorithm with the following priors: δ_*ik*_ ∼ *IG*(0.5,0.5) and Σ ∼ *IW*(5,*I*_50_). A chain of 15,000 iterations was generated and the last 10,000 samples were kept for calculating the posterior mean.

Figures 1 shows the estimation bias, i.e., *P̂*(*y*\* = 1 |***x***\*) – *P*(*y*\* = 1 |***x***\*), by two models under two settings. In both settings, the copula-based method shows its superiority over the independence model, and the improvement is more significant in the presence of stronger correlation. Since moderate and strong coexpression between genes were commonly seen in real data, the ignorance of such information may lead to lower prediction accuracy.

**Figure 1.**
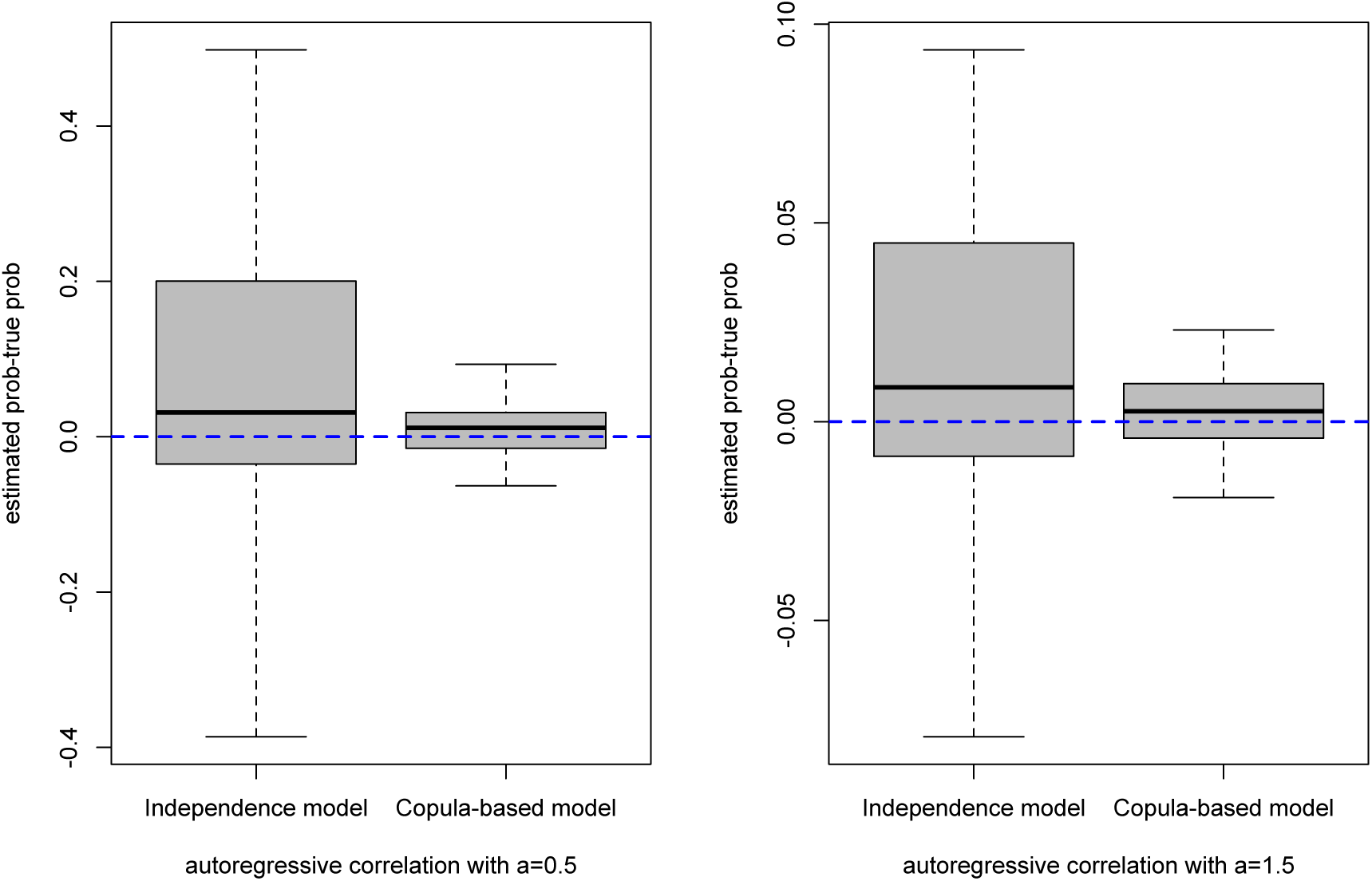
Comparison between Dong et al.’s independence model and the copula-based model. The y-axis represents the estimation bias in the discriminant score, i.e., the estimated score minus the true score.

### 3.2 Simulation II

In the second simulation, we compared our copula-based classifier with the other five classifiers including Poisson linear discriminant analysis (PLDA), negative binomial linear discriminant analysis (NBLDA), k-nearest neighbors (KNN), partial least square method (PLS) and logistic regression method. We implemented PLDA using R package *PoiClaClu* and implemented NBLDA using the R source code provided by the Dong et al. (https://github.com/yangchadam/NBLDA). In NBLDA, the overdispersion parameters were estimated by R package *sSeq*. For KNN, we chose parameter *K* = 1,3,5. To implement PLS, we used the function ’spls()’ provided in R package *spls.* For our new classifier, same priors were used as in Simulation I. A chain of 15,000 iterations was generated and the last 10,000 samples are retained for estimation.

The data are generated under four different settings (for all settings, *μ_i_*_1_ = 20, *μ_i_*_2_ = *μ_i_*_1_ + Δ_*i*_, Δ_*i*_ ∼ *Unif* (–15,15)):

- Setting 1 (weaker correlation, smaller dispersion): Ω_*k*_(*i*, *j*) = exp(–1.5|*i* – *j* |), δ_*ik*_ ∼ *Unif*(1,5), *k* = 1,2
- Setting 2 (weaker correlation, larger dispersion): Ω_*k*_(*i*, *j*) = exp(–1.5|*i* – *j* |), δ_*ik*_ ∼ *Unif*(5,20), *k* = 1,2
- Setting 3 (stronger correlation, smaller dispersion): Ω_*k*_(*i*, *j*) = exp(–0.5|*i* – *j* |), δ_*ik*_ ∼ *Unif*(1,5), *k* = 1,2
- Setting 4 (stronger correlation, larger dispersion): Ω_*k*_(*i*, *j*) = exp(–0.5|*i* – *j* |), δ_*ik*_ ∼ *Unif*(5,20), *k* = 1,2

The comparison results under different sample sizes (*n* = 10,30,60,100) are shown in Figures 2 and 3. Due to the independence assumption, other five classifiers including PLDA and NBLDA failed to model the dependence structure between genes, therefore the estimated probabilities were biased (see Simulation I). It is observed that the copula-based model performs consistently better than the other classifiers in terms of classification accuracy, especially in setting 4 with stronger correlation and larger dispersion. When the correlation between genes are very weak, our model has similar performance with Dong et al.’s independence model.

**Figure 2.**
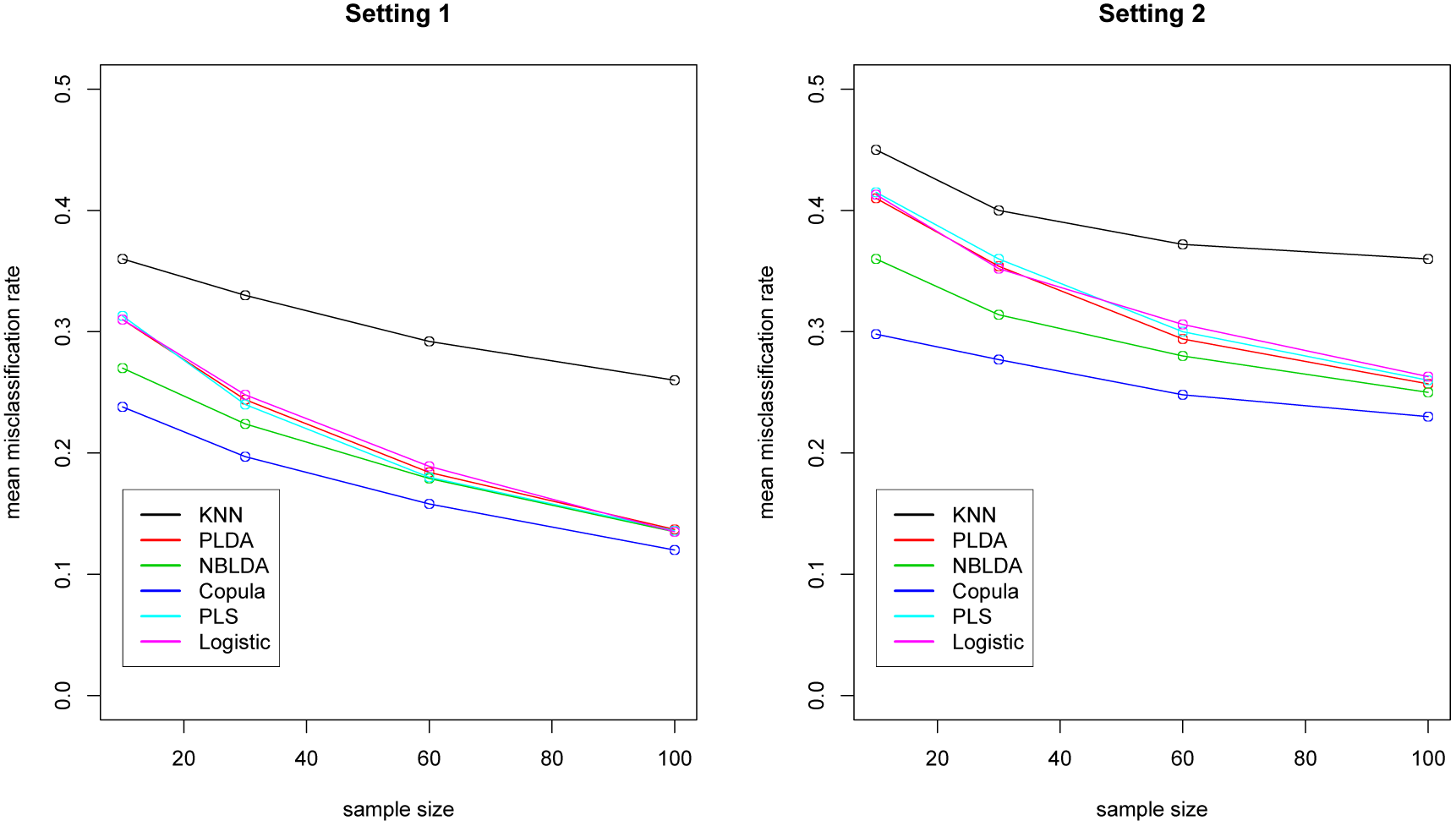
Comparison of six different classifiers under setting 1 (weaker correlation and smaller dispersion) and setting 2 (weaker correlation and larger dispersion). The y-axis is mean misclassification rate based on 100 independent simulation runs. The x-axis is sample size, *n* = 10,30,60,100.

**Figure 3.**
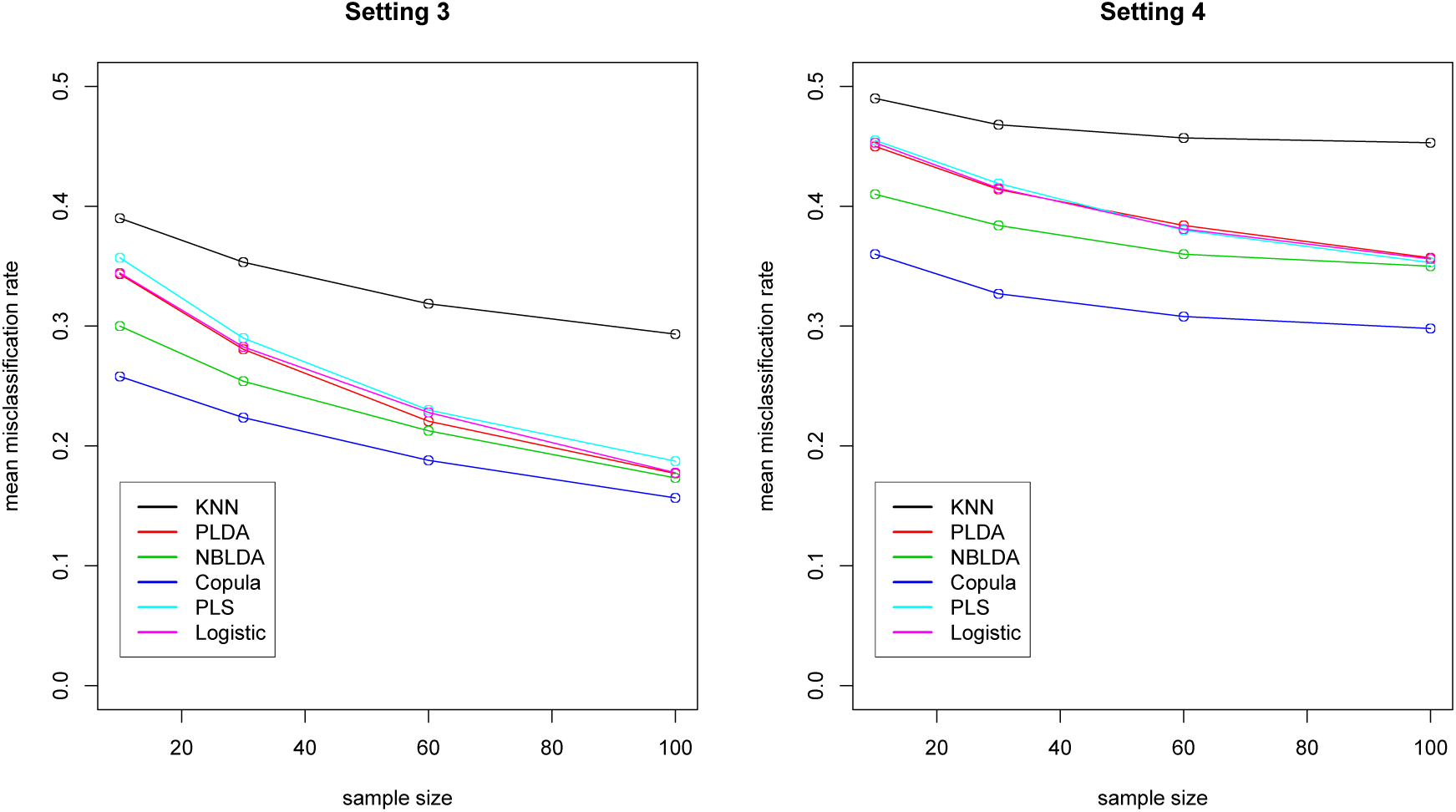
Comparison of six different classifiers under setting 3 (stronger correlation and smaller dispersion) and setting 4 (stronger correlation and larger dispersion). The y-axis is mean misclassification rate based on 100 independent simulation runs. The x-axis is sample size, *n* = 10,30,60,100.

## 4 Real data analysis

In this experiment, we considered two real data sets including the cervical cancer data (Witten et al. [2010], available in Gene Expression Omnibus (GEO) with access number GSE20592) and the HapMap data (Montgomery et al. [2010]; Pickrell et al. [2010], available at ftp://ftp.ncbi.nlm.nih.gov/hapmap). The cervical cancer data contains 58 samples (29 tumor samples and 29 normal controls), and 714 microRNAs which were differentially expressed in cancer group and normal group. The HapMap data contains 129 samples (60 CEU samples and 69 YRI samples) and a total number of 52,580 genes. For both data sets, we removed genes with less than 10 reads across all samples. Three different classifiers including Poisson classifier, negative binomial classifier and the copula-based classifier were compared in terms of the misclassification rate. In our classifier, same priors were used as in the simulation studies.

We noted that the real data sets often contain large portion of irrelevant and redundant genes. A gene screening could greatly reduce the computing time and improve the classification accuracy. We conducted gene selections using R package *edgeR* (available in *Bioconductor,* www.bioconductor.org), as suggested by Dong et al. The algorithm implemented in *edgeR* is based on negative binomial model and takes overdispersion into account, therefore it is suitable for our problem. This method first estimates the overdispersion parameter for each gene by maximizing the combination of gene-specific conditional likelihood and the overall conditional likelihood, and then constructs an exact test using negative binomial distribution.

For Cervical cancer data, 40 samples were randomly assigned to the training set and the rest 18 samples were assigned to the testing set. A total of 20, 50, 100, 300 genes were selected, respectively. For HapMap data, the samples were randomly split into training set and testing set, with 70 samples and 59 samples, respectively. A total of 50, 100, 300, 500 genes were selected, respectively. Three different classifiers were then trained by the training data and applied to the testing data. The whole procedure were repeated for 100 times and the average misclassification rate were recorded.

The comparison results are shown in Figure 4. For both data sets, the copula-based model is more accurate than the other two classifiers. Figure 5 displays the distribution of correlation coefficients between every pair of genes in the cervical cancer data (log-transformed). It can be seen that the vast majority of the correlations are positive, and half of them are fairly strong (above 0.5), indicating that the independence assumption in PLDA and NBLDA is violated.

**Figure 4.**
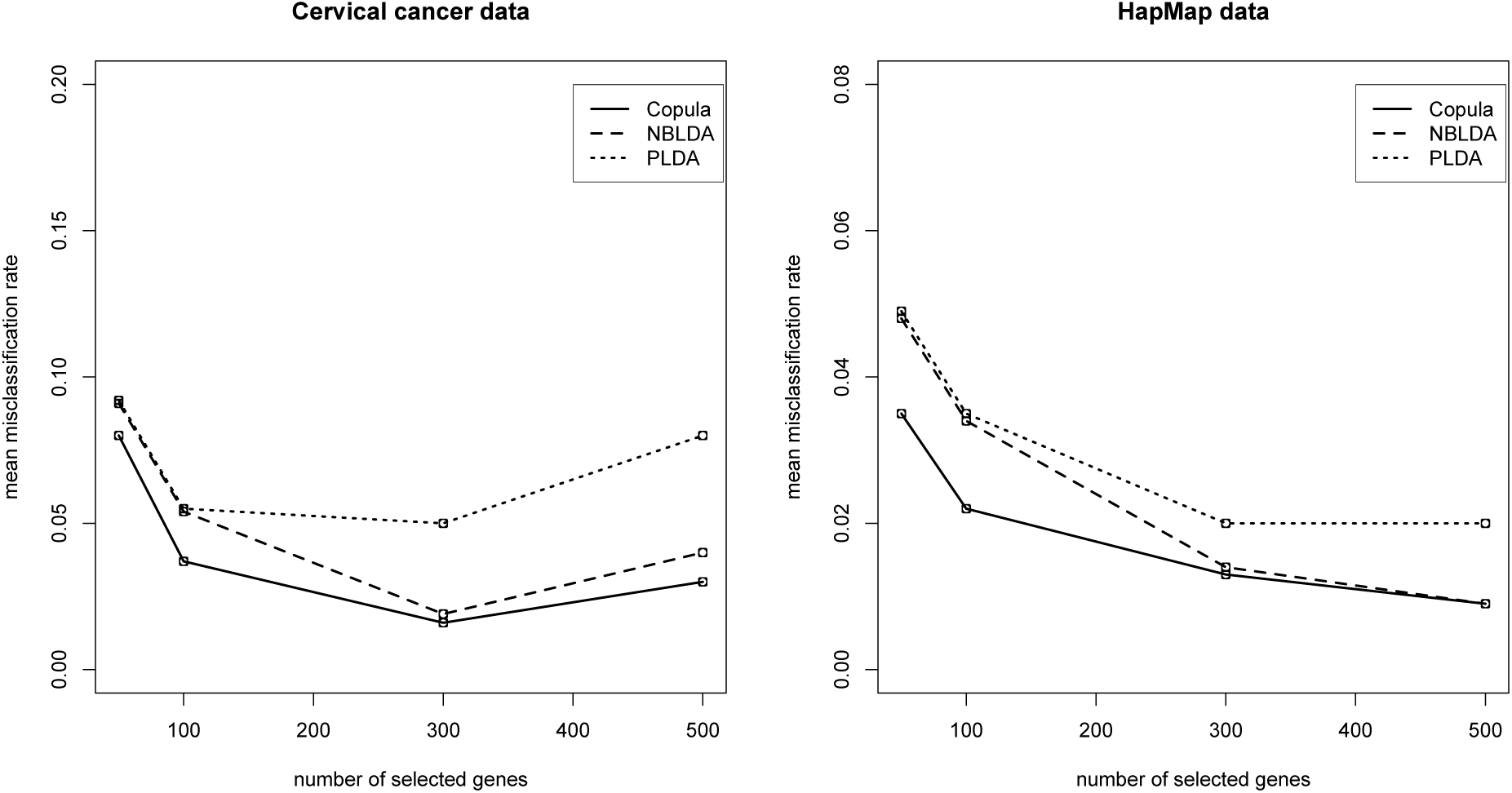
Comparison of three classifiers on two real data sets: cervical cancer data and HapMap data. The y-axis is mean misclassihcation rate based on 100 runs. The x-axis is number of selected genes, *p* = 50,100,300,500

**Figure 5.**
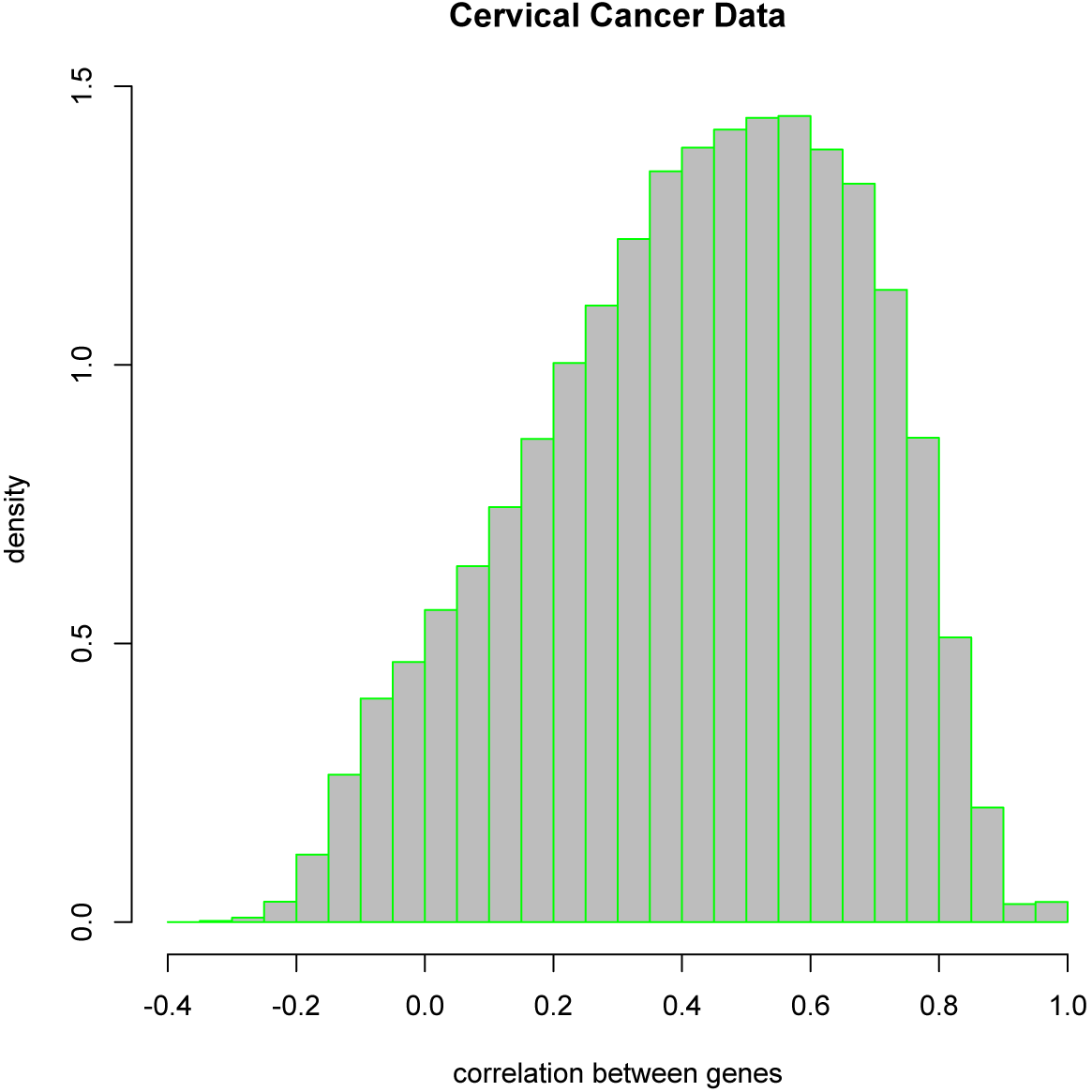
Distribution of correlations between every pair of genes in the cervical cancer data. The Pearson’s correlation coefficients are based on the log-transformed data. About 93.49% of the gene pairs show positive correlation and only 6.51% of gene pairs show negative correlation.

## 5 Discussion

In this paper, we have proposed a new classifier for RNA-Seq data. Different from other classifiers, it incorporates the dependence between genes in the supervised classification problem. To the best of our knowledge, this is the first work that applies copula model to the classification of count data. Numerical comparisons show that our new model achieves better estimate of discriminant scores than existing methods, therefore results in more accurate sample classification. The improvement is more significant in the presence of stronger correlation between genes. In addition to RNA-Seq data, this classifier can be generally applied to other digital gene expression data to improve the classification accuracy.

The copula-based classifier introduced in this paper assumes that the reads count of each gene follows a negative binomial distribution. This assumption has been widely used in practice since the negative binomial distribution is flexible to model and quantify the overdispersion in RNA-seq data. However, the negative binomial assumption still might be violated in some real data sets. In the copula-based method, one could choose alternative marginal models (e.g., Poisson mixture model) which better fits the data, and the Bayesian estimation introduced in this paper still can be applied to estimate the covariance matrix since the estimation of copula only depends on the cumulative distribution functions of the marginals. It is also noteworthy that the current PXMH algorithm for parameter estimation is time-consuming when the number of selected genes is large. For example, in the analysis of HapMap data, a single run (out of 100 runs in total) with 500 selected genes takes about 2.5-3 minutes with C++ implementation. In the future study, we would like to explore other optimizations such as EM-type method for computational efficiency. For example, rather than joint estimating {δ, Ω} in Bayesian framework, we may first estimate δ using stable method based on read counts, and then estimate Ω by EM update where latent variables **Z** can be treated as missing values. The EM-type method may greatly reduce the computing time. To model the correlation between genes, we use Gaussian copula as it is convenient for multivariate problem. Another possible future work is to compare different latent variables and copula functions, e.g., Student’s t copula (Nelson [1999]) and Gaussian mixture copula (Zhang & Shi [2016]), in a model comparison framework (Mai & Zhang [2016], Zhang et al. [2014],

Matveeva et al. [2016]).

## 6 Conclusion

RNA-sequencing experiment quantify gene expression by the count of short reads mapped to the gene region. When biological replicates are available, the negative binomial distribution allowing overdispersion is better suited for modeling RNA-Seq data than Poisson distribution. Recently, Dong et al. (Dong et al. [2016]) developed a classifier based on negative binomial distribution, which outperforms previous methods including the Poisson classifier and K-nearest neighbors classifier. However, due to the difficulty of modeling the dependence in discrete data, most existing classifiers assume that all the genes are independent of each other. In this paper, we systematically investigate the effect of independence assumption on discriminant score calculation and classification. In addition, we developed a copula-based classifier for RNA-Seq data that incorporates the dependence structure between genes, while maintaining the negative binomial marginals. Our numerical comparisons and real data analysis demonstrate that the new classifier performs better than existing methods including Dong et al.’s negative binomial classifier.

## Acknowledgement

Support has been provided in part by the Arkansas Biosciences Institute, the major research component of the Arkansas Tobacco Settlement Proceeds Act of 2000.

## Abbreviations

PLDA: : Poisson linear discriminant analysis;
NBLDA: : Negative binomial linear discriminant analysis;
KNN: : K-nearest neighbors;
PXMH: : Parameter-expanded reparameterization and Metropolis-Hasting

## Appendix

### Sampling *z_ij_* conditioning on {*z*_–*i*,*j*_, δ_*i*_, Ω}

We sample *z_ij_* from univariate normal distribution ϕ(*Z_ij_*|*μ_ij_*,*ω_ij_*^2^) truncated between *L_ij_·* and *U_ij_*:

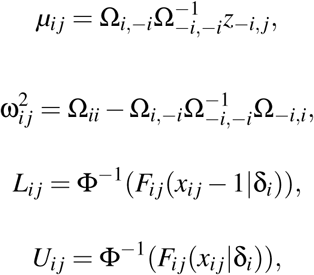

where Ω *_i,–i_ =* {Ω*_hl_*, *h* = *i, l* ≠ *i*}, Ω_−*i,i*_ = {Ω*_hl_,h* ≠ *i, l* = *i*}, Ω_*−i,−i*_ = {Ω_*hl*_, *h* ≠ *i, l* ≠ *i*} and *z*_−i,j_ = (*z*_1 *j*_, …, *z*_(*i*−1)*j*_, *z*_(*i*+1)*j*_, …, *z_pj_*).

### Sampling δ*_i_* conditioning on {*z*_–*i*_,·, δ_–*i*_, Ω}

We sample δ_*i*_ based on the following density function:

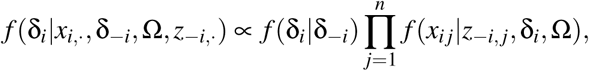

Where 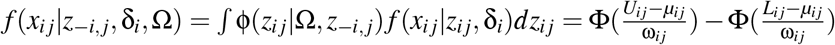.

Suppose δ_*i*_ is the current value and δ_*i*_* is the generated value from the proposal distribution. The Metropolis-Hasting probability of moving from δ_*i*_ to δ_*i*_* is:

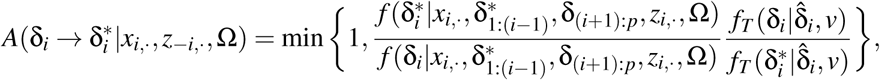

where the proposal *f_T_*(δ_*i*_|δ̂_*5*_,*ν*) is a t distribution which dominates the normal tails, δ̂_*i*_ and *ν* represents location and degree of freedom, respectively.

### Sampling Ω conditioning on {*z*, δ}

Sampling of Ω can be problematic due to the constraint Ω*_ii_*= 1,*i =* 1,…,*p.* Here we sample Ω using parameter-expanded reparameterization and Metropolis-Hasting algorithm (PXMH, Lee [2014]; Liu & Daniels [2006]). The conditional density can be written as follows:

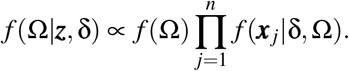

The PXMH algorithm first simulates a covariance matrix Σ and then transforms it to a correlation matrix Ω. For convenience, define 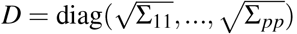, then Ω = D^−1^ΣD^−1^. Since Ω has *p* fewer parameters than Σ, an additional constraint is imposed:

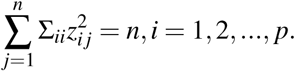

The PXMH Algorithm consists of three steps:

- PX step: Sample Σ ∼ *IW_p_*(*ν*,Ψ), where *ν* = *ν*_0_ + *n* and 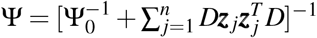
- MH step: Move to the new value Σ* with probability:

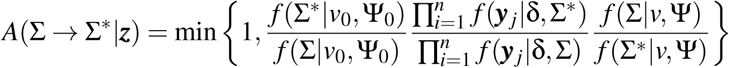
- Transform covariance matrix Σ to correlation matrix Ω, Ω* = *D*\*^−1^ Σ* *D*\*^−1^

